# Quorum-Sensing Stimulation and Phytochemical Quenching Reshape Biofilm-Associated Gene Expression in *Salmonella enterica*

**DOI:** 10.64898/2026.05.26.727871

**Authors:** Schnieder Fernandes, Anjali Ghosh, Caroline Smith, Ihab Tewfik, Kalpana Surendranath, Vincenzo Torraca, Genomics and Infectious Disease Research Group

## Abstract

Quorum sensing (QS) influences biofilm formation, persistence and stress adaptation in *Salmonella enterica*. Although *Salmonella* does not synthesise acyl-homoserine lactones (AHLs), it can detect exogenous AHLs through the LuxR homolog SdiA, allowing it to respond to interspecies signalling cues in polymicrobial environments. This study investigated whether external QS stimulation and quorum-modulatory compounds reshape biofilm-associated transcriptional programmes in *S. enterica* serovar Enteritidis (SE) and *S. Typhimurium* ST14028. Biofilm formation was assessed using the crystal violet assay, while expression of QS-, biofilm-, adhesion-, motility- and invasion-associated genes (*sdiA, csgD, flgG, fimA, rck, invA, bapA* and *hilA*) was quantified using multiplex RT-qPCR and analysed by the ΔΔCt method, with 16S rRNA used for normalisation. In parallel, molecular docking was used to explore the predicted interaction of C8-HSL, established quorum-quenching agents and selected phytochemicals with the *Salmonella* SdiA ligand-binding region. Exposure to exogenous C8-HSL increased biofilm biomass and induced coordinated upregulation of QS- and biofilm-associated genes in both serovars, supporting the role of external AHL sensing in *Salmonella* biofilm regulation. In contrast, farnesol and furanone produced broad transcriptional repression accompanied by reduced biofilm biomass. Selected natural products, including epigallocatechin gallate (EGCG), thymoquinone, garlic extract, turmeric extract and aloe-emodin, produced moderate antibiofilm effects and partial downregulation of QS-associated transcriptional responses, suggesting possible interference with biofilm-regulatory signalling pathways. Molecular docking further supported this interpretation by identifying potential interactions between selected quorum-modulatory compounds and the predicted SdiA binding region, providing a plausible mechanistic basis for their observed biological effects. Notably, responses differed between SE and ST14028, indicating strain-dependent sensitivity to QS stimulation and quorum-modulatory treatments. Collectively, these findings suggest that exogenous AHL sensing contributes to strain-dependent transcriptional reprogramming of *Salmonella* biofilm-associated genes and that selected phytochemicals may act as preliminary quorum-modulatory candidates. This study supports further investigation of SdiA-mediated signalling as an anti-virulence target for reducing *Salmonella* persistence in food-associated and clinical environments.

## 1. Introduction

*Salmonella* enterica remains one of the leading causes of foodborne disease worldwide, responsible for ∼93 million cases of gastroenteritis and approximately 155,000 deaths each year (Lamichhane et al., 2024). In the United Kingdom, *Salmonella* continues to represent a significant and resurging public health concern. Recent UK Health Security Agency (UKHSA) surveillance reported 10,388 laboratory-confirmed non-typhoidal *Salmonella* infections in England in 2024, representing a 17.1% increase from 2023 and the highest level recorded in the past decade. Notably, children under 10 years accounted for over one-fifth of cases, highlighting the ongoing vulnerability of key population groups (Lamichhane et al., 2024). Beyond community incidence, the clinical burden is reflected in healthcare utilization, with 1,468 hospital admissions for salmonellosis recorded in England between April 2022 and March 2023 (approximately 3 per 100,000 population), representing a substantial increase compared with previous years recorded by the NHS. *Salmonella* also remains a major contributor to foodborne outbreaks in the UK, accounting for approximately 27% of reported foodborne outbreaks between 2015 and 2020 reported by food standards agency UK. The persistence of S. enterica within food chains and clinical environments, coupled with its capacity to cause both acute gastroenteritis and prolonged carriage, continues to pose major challenges for infection control, food safety, and clinical management. These epidemiological trends underscore the urgent need for alternative control strategies that target key pathogenic mechanisms beyond conventional antimicrobial approaches. (Zhang et al 2025)

The remarkable persistence and epidemiological success of *Salmonella* enterica are largely driven by its considerable genetic adaptability, which enables rapid adjustment to diverse host and environmental conditions (Sholpan et al., 2021, Mi et al., 2024). This Gram-negative pathogen coordinates a complex regulatory network controlling adhesion, invasion, stress tolerance, and—critically—biofilm formation, a trait strongly associated with chronic contamination and reduced antimicrobial susceptibility (Singh et al., 2025; Małaszczuk et al., 2025). The continued global rise in antimicrobial resistance among *Salmonella* strains has further complicated treatment and control efforts, highlighting an urgent need for alternative strategies that do not rely solely on conventional bactericidal approaches (Hetta et al., 2024; Lamichhane et al., 2024).

In this context, quorum sensing (QS) has emerged as a promising anti-microbial target (Chu et al., 2024). By modulating bacterial communication pathways rather than directly inhibiting growth, QS interference offers the potential to attenuate pathogenic behaviors such as biofilm formation while exerting reduced selective pressure for resistance development (Zhao et al., 2025; Hetta et al., 2024). Accordingly, the present study focuses on characterizing quorum sensing–mediated biofilm regulation in *Salmonella* enterica, with the aim of evaluating whether targeting QS pathways may provide a viable strategy to mitigate persistence and pathogenicity in this clinically significant organism. Biofilm formation is a central determinant of *Salmonella* enterica persistence in both clinical and food-associated environments (Lamichhane et al., 2024) (Małaszczuk et al., 2025). The biofilm extracellular polymeric substance (EPS) matrix—composed primarily of curli fimbriae, cellulose, and large surface adhesins—confers enhanced tolerance to antimicrobial agents, environmental stress, and host immune clearance (Xiang et al., 2024; Małaszczuk et al., 2025). At the regulatory level, the transcription factor CsgD functions as the master controller of biofilm development, coordinating the expression of key structural components including BapA and fimbrial systems (Sokaribo et al., 2020; Chen et al., 2021).In addition to matrix production, successful biofilm establishment requires coordinated regulation of adhesion, motility, and invasion-associated genes. Factors such as FimA, Rck, HilA, InvA, and the flagellar component FlgG contribute to early surface attachment, host interaction, and biofilm maturation, highlighting the multi-gene nature of *Salmonella* biofilm regulation. (Zhang et al., 2022; Sholpan et al., 2021)

Importantly, these biofilm-associated pathways are modulated by quorum sensing (QS), a cell-density–dependent communication mechanism that enables bacteria to synchronise group behaviours. Although S. enterica does not synthesise acyl-homoserine lactones (AHLs), it possesses the LuxR homolog SdiA, which enables detection of exogenous AHL signals produced by neighbouring microbial species. This interspecies sensing capability allows *Salmonella* to exploit environmental signalling cues to enhance virulence and biofilm formation (Sholpan et al., 2021; Zhang et al., 2022; Lamichhane et al., 2024). Experimental studies have shown that supplementation with medium-chain AHLs, particularly C8-HSL, can upregulate QS-responsive genes and promote biofilm development in *Salmonella*. This makes the SdiA–AHL signalling axis an attractive anti-virulence target, particularly in the context of rising antimicrobial resistance where strategies that attenuate pathogenic behaviour without inhibiting growth are increasingly sought (Dyszel et al., 2010; Zhang et al., 2022; Hetta et al., 2024).Consequently, there is growing interest in natural products as potential quorum sensing modulators. Phytochemicals such as polyphenols, quinones, and plant-derived extracts have demonstrated the ability to interfere with QS signalling and biofilm formation in Gram-negative pathogens (Fleitas Martínez et al., 2019; Cheng et al., 2024). However, their transcriptional impact on key QS-biofilm genes in clinically relevant *Salmonella* strains remains insufficiently characterised. Accordingly, the present study investigates the effect of selected natural compounds on QS-mediated biofilm regulation in *Salmonella enterica*. Using exogenous C8-HSL to stimulate quorum sensing, we quantified both phenotypic biofilm formation and transcriptional changes in key regulatory and structural genes (sdiA, csgD, bapA, fimA, rck, hilA, invA, flgG). This integrated approach aims to determine whether natural products can meaningfully disrupt quorum-regulated biofilm pathways and thereby represent viable anti-virulence candidates.

Given the central role of quorum sensing in coordinating *Salmonella* biofilm formation, increasing attention has focused on naturally derived compounds capable of interfering with bacterial communication pathways (Shamim et al., 2023; Alum et al., 2025). Natural products are particularly attractive as quorum sensing modulators because many possess multi-target activities, relatively low toxicity profiles, and reduced propensity to drive classical antimicrobial resistance ((Hetta et al., 2024; Alum et al., 2025; Defoirdt, 2025). In the present study, a targeted panel of phytochemical and biologically derived agents was selected based on reported or proposed quorum sensing and antibiofilm activity in Gram-negative pathogens. Epigallocatechin gallate (EGCG), a major green tea polyphenol, has been shown to disrupt AHL-mediated signalling and downregulate QS-regulated virulence genes in enteric bacteria. Aloe-emodin, an anthraquinone derivative, has demonstrated interference with bacterial communication systems and inhibition of biofilm maturation in several Gram-negative species (Cheng et al., 2024; Hao et al., 2021; Chi et al., 2022). Similarly, thymoquinone, the principal bioactive component of *Nigella sativa*, has been reported to attenuate quorum sensing–controlled phenotypes and reduce biofilm biomass (Dera et al., 2021; Hetta et al., 2024)

Complex natural extracts were also included to capture multi-component quorum-modulatory effects. Garlic extract contains organosulfur compounds such as allicin that have been widely associated with quorum quenching activity and disruption of bacterial signalling networks s (Morshdy et al., 2022; Shamim et al., 2023). Turmeric extract, rich in curcuminoids, has been reported to interfere with biofilm regulatory pathways and surface adhesion mechanisms. Commercial honey provides a multifactorial antimicrobial environment combining osmotic stress, hydrogen peroxide generation, and phytochemical content, while cinnamon-derived compounds have shown potential to interfere with AHL signalling and bacterial surface attachment. (Chi et al., 2022; Matei et al., 2025; Aswathanarayan et al 2018) To benchmark these natural compounds against established quorum modulators, we included C8-HSL as a positive QS inducer and the well-characterised quorum quenchers farnesol and furanone as inhibitory controls, alongside chlorhexidine and untreated LB controls. This structured panel enables discrimination between general growth inhibition and specific quorum-mediated antibiofilm effects (Zhang et al., 2022; Tan et al., 2024; Hetta et al., 2024). By integrating phenotypic biofilm quantification with RT-qPCR analysis of key QS-responsive genes, this study aims to determine whether these natural products exert measurable quorum-modulatory effects in *Salmonella enterica* and to identify candidates with potential anti-virulence utility.

## 2. Materials and Methods

### 2.1 Bacterial Strains and Culture Conditions

*Salmonella enterica* serovar Enteritidis UCH (University of Westminster culture collection), *Salmonella Typhimurium* ATCC 14028, and *Escherichia coli* K-12 MG1655 were obtained from the University of Westminster microbiology repository. Where applicable, strains were originally sourced from the American Type Culture Collection (ATCC). Bacterial stocks were maintained on Luria–Bertani (LB) agar plates at 37 °C supplemented with ampicillin (100 µg/mL) for E.coli and Kanamycin (50ug/ml) for *Salmonella* where required. Isolates were confirmed using the API 20E identification system (bioMérieux, France), and all *Salmonella* serovars were subjected to antimicrobial susceptibility testing.

For experimental assays, single colonies were inoculated into LB broth and incubated overnight at 37 °C with shaking at 150 rpm using an INNOVA42 shaking incubator (Eppendorf). Overnight cultures were standardised to an optical density (OD₅₇₀) of 1.0 (approximately 1 × 10⁷ CFU/mL) and used as the starting inoculum for downstream experiments. All experiments were performed using independent biological replicates under identical growth conditions. Work involving *Salmonella* was conducted in accordance with institutional biosafety guidelines.

### 2.2 Quorum sensing, quorum quenching and natural product molecules

#### 2.2.1 Quorum sensing and quorum quenching reference molecules

*N-(3-oxooctanoyl)-L-homoserine lactone* (C8-HSL) was purchased from Cayman Chemical Company (USA). Farnesol and halogenated furanone were obtained from Sigma-Aldrich (Merck, UK) and were used as established quorum-quenching controls.

For each QS-active or quorum-quenching molecule, a 1 mM stock solution was prepared in ethyl acetate, filter-sterilised (0.22 µm), aliquoted, and stored at −20 °C in the dark until use. Working concentrations of 0.20 mM, 0.05 mM, and 0.01 mM were freshly prepared prior to experiments by serial dilution of the 1 mM stock into sterile LB broth. The final ethyl acetate concentration in all cultures, including solvent controls, did not exceed 1% (v/v), and an ethyl acetate–only control was included in all assays. All other chemicals used were of analytical grade. Prior to experiments 0.2mM was identified to be the appropriate concentrations of all experimental groups through a series of MIC and growth assays.

#### 2.2.2 Natural product preparations

All natural product solutions or suspensions were freshly prepared on the day of use, unless otherwise stated, and were passed through a 0.22 µm filter where feasible to minimise microbial contamination.

##### Garlic extract

Fresh garlic cloves (Allium sativum) were peeled, weighed, lyophilized, and then homogenised in sterile phosphate-buffered saline (PBS; pH 7.4) at a ratio of 1 g tissue per 5 mL PBS using a sterile mortar and pestle or homogeniser. The homogenate was incubated at room temperature for 30 min to allow allicin formation, then centrifuged at 5,000 × g for 10 min at 4 °C. The supernatant was collected, passed through a 0.22 µm syringe filter, and used as crude garlic extract. An appropriate volume of sterile PBS was used to obtain the desired working concentration in LB broth (100mg/ml), and PBS-only controls were included.

##### Turmeric extract

Dried turmeric (Curcuma longa) powder was weighed, lyophilized and extracted in 70% (v/v) ethanol at a ratio of 1 g powder per 10 mL solvent, followed by vortexing and incubation at room temperature for 1 h with occasional agitation. The mixture was then centrifuged at 5,000 × g for 10 min, and the supernatant was collected and filtered (0.22 µm). The ethanolic extract was used to prepare a stock solution (100 mg/mL), stored at −20 °C protected from light, and diluted into LB broth to the desired final concentration immediately before use. Corresponding ethanol-only controls (same final % v/v) were included to control for solvent effects.

##### Epigallocatechin gallate (EGCG)

Epigallocatechin gallate (EGCG; ≥95% purity; Sigma-Aldrich, UK; Cat. No. E4268) was used as a natural quorum sensing and biofilm modulator. A 10 mM stock solution was prepared in sterile distilled water, filter-sterilised (0.22 µm), aliquoted, and stored at −20 °C protected from light. Working solutions were freshly prepared in LB broth at the required final concentrations immediately prior to inoculation.

##### Aloe-emodin

Aloe-emodin (analytical grade; Sigma-Aldrich, UK; Cat. No. A7687) was included as a natural QS/biofilm modulator. Due to its limited aqueous solubility, a concentrated stock solution (20 mM) was prepared in dimethyl sulfoxide (DMSO), vortexed until fully dissolved, filter-sterilised (0.22 µm), aliquoted, and stored at −20 °C protected from light. Prior to use, the stock was diluted into LB broth to achieve the desired final concentration, ensuring that the final DMSO concentration was matched in solvent control wells.

##### Thymoquinone

Thymoquinone (analytical grade; Sigma-Aldrich, UK; Cat. No. 274666) was used as a natural quorum sensing modulator. A concentrated stock solution (50 mM) was prepared in DMSO, vortexed thoroughly until fully dissolved, aliquoted, and stored at −20 °C to minimise freeze–thaw cycles. Immediately prior to experiments, the stock was diluted into LB broth to the required final concentrations, with equivalent DMSO levels maintained in all corresponding control wells.

#### 2.2.3 Biocidal and antimicrobial control

Chlorhexidine digluconate solution (Sigma-Aldrich, UK; Cat. No. C9394) was used as a non-QS-specific biocidal control. The supplied aqueous stock solution was used to prepare sterile working solutions in LB broth immediately prior to inoculation. Where required, the solution was filter-sterilised (0.22 µm). Sterile distilled water controls were included in all corresponding assays. Chloramphenicol was used as antimicrobial control group as the Salmonella isolates showed inhibition at 16ug/ml.

#### 2.2.4 Final treatment setup

For all experiments, treatment molecules were added to LB broth to achieve the desired final concentrations prior to inoculation with *Salmonella* cultures. C8-HSL was tested at 0.20 mM; farnesol at 0.20 mM; halogenated furanone at 0.10 mM; EGCG at 0.20 mM; aloe-emodin and thymoquinone at 0.10 mM; garlic and turmeric extracts at 0.5–2% (v/v); ; cinnamon preparations at 0.25–1%; and chlorhexidine digluconate at 0.002–0.02% (w/v). Untreated LB broth and appropriate solvent controls (DMSO, ethanol, PBS, or water at matched final concentrations) were included in parallel. All treatments were prepared in at least triplicate wells for growth curve, biofilm, and RT-qPCR assays.

### 2.3 CV assay biofilm quantification

Following the final OD measurement at 72 hours, all 12-well plates were subjected to a crystal violet (CV) staining assay to quantify biofilm formation. Plates were gently shaken over a waste container to discard planktonic (free-floating) cells, preserving only the adherent cells attached to the well surface, representing biofilm biomass.

Each well was then rinsed thoroughly with distilled water to remove remaining planktonic cells. At this stage, a portion of the biofilm was collected and stored for subsequent DNA and RNA extraction. To stain the biofilm, 500 µL of 0.1% crystal violet solution was added to each well and incubated for 15 minutes at room temperature. After staining, wells were rinsed with distilled water to remove excess dye, ensuring uniform staining of biofilm without overstaining.

The plates were then incubated at 60 °C for 1 hour to remove residual moisture. Once dry, 250 µL of 33% acetic acid was added to each well to solubilize the bound dye. The plates were incubated for 10 minutes, after which the contents of each well were mixed thoroughly by pipetting up and down. The solubilized dye was transferred to a fresh 12-well plate to eliminate background interference from residual dye or biomass. Absorbance was measured at 570 nm using a microplate reader to quantify biofilm biomass.

### 2.4 Polymerase Chain Reaction (PCR) and Reverse Transcription PCR (RT-PCR)

#### 2.4.1 DNA and RNA Extraction

##### DNA Extraction via the Boiling Method

Bacterial pellets from a biofilm culture were resuspended in RNase-free water and subjected to heat treatment at 100 °C for 15 minutes in a heat block to lyse the cells and release genomic DNA. Immediately after heating, the tubes were cooled on ice to minimise DNA degradation. The samples were then centrifuged at maximum speed, and the supernatant—containing the extracted DNA—was carefully transferred to fresh, labelled 1.5 mL microcentrifuge tubes. The DNA was used for endpoint PCR.

##### RNA Extraction

Bacterial cultures were grown in LB broth for 8 hours in the presence of the respective treatments. Growth was halted based on growth curve data indicating that cells had reached the late exponential to early stationary phase at this time point. Total RNA was extracted using the SPINeasy RNA Kit for Bacteria with Lysozyme (MP Biomedicals, UK) according to the manufacturer’s instructions.

Briefly, 1.5 mL of bacterial culture was harvested by centrifugation and the supernatant discarded. The cell pellet was subjected to enzymatic lysis using the kit-provided lysozyme buffer, followed by binding of RNA to the silica membrane column. After sequential wash steps to remove contaminants and genomic DNA, RNA was eluted in nuclease-free water. RNA samples were either used immediately for one-step RT-PCR or stored at −20 °C for short-term use and −70 °C for long-term storage.

#### 2.4.2 Primer Selection and Endpoint PCR

Primers specific to *Salmonella* spp. are listed in Table 1 below. Primer specificity was verified using NCBI-BLAST2 via the EMBL-EBI platform.

**Table 1.**
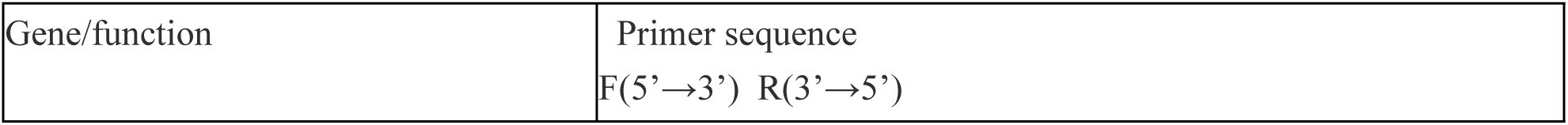
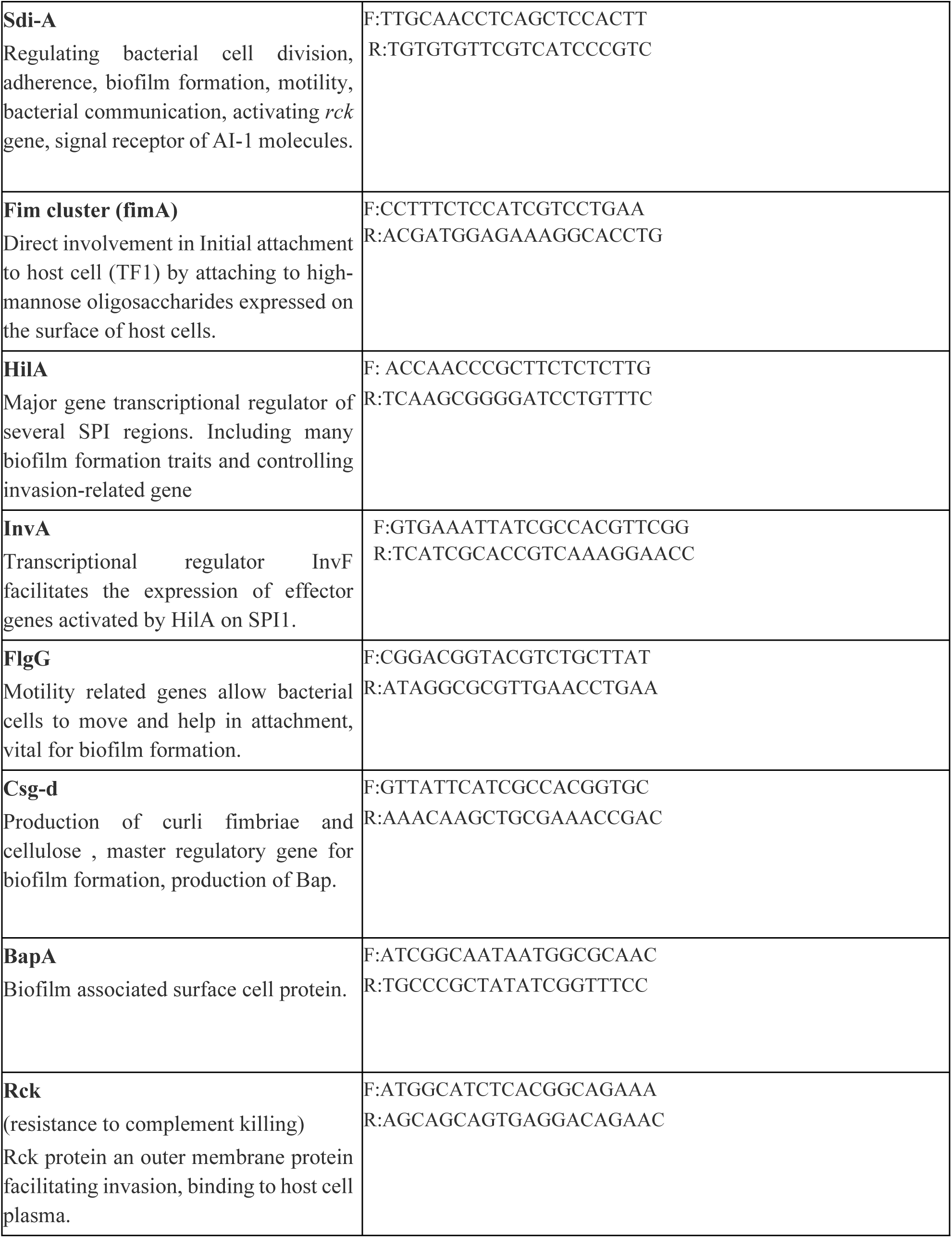

Each PCR reaction was prepared in a total volume of 25 µL, consisting of 12.5 µL of Master Mix, 2.5 µL of forward primer, 2.5 µL of reverse primer, 2 µL of template DNA, and 4.4 µL of sterile distilled water. Thermal cycling conditions included an initial denaturation step at 94 °C for 3 minutes, followed by 35 cycles of denaturation at 95 °C for 30 seconds, annealing at 55 °C for 30 seconds, and extension at 72 °C for 30 seconds. A final extension was carried out at 72 °C for 5 minutes. PCR products (5 µL) were analysed on 1% agarose gels prepared in 1× TAE buffer containing ethidium bromide (0.5 µg/mL) and electrophoresed at 135 V for 2 hours. A 100 bp DNA ladder was included as a molecular weight marker.

#### 2.4.3 Reverse Transcription Quantitative PCR (RT-qPCR)

Multiplex RT-qPCR was performed using gene-panel assays designed to simultaneously quantify quorum-sensing and biofilm-associated genes in *Salmonella* Enteritidis, *Salmonella* Typhimurium ST14028 and *E. coli*. For *Salmonella*, three multiplex panels were used: P1 (sdiA–CY5, csgD–Texas Red, flgG–HEX, 16S–FAM), P2 (fimA–CY5, rck–Texas Red, invA–HEX, 16S–FAM), and P3 (bapA–CY5, hilA–Texas Red, 16S–FAM). For *E. coli,* two panels were used: P1E (sdiA, csgD, flgG, 16S) and P2E (fimA, 16S). RNA was extracted from cultures grown under all treatment conditions (LB, C8-HSL, Farnesol, Furanone, Chlorhexidine, EGCG, thymoquinone, Garlic, Turmeric and Aloe-emodin), treated with DNase I, and converted to cDNA using 500 ng–1 µg RNA per reaction. qPCR reactions were assembled in 25 µL volumes containing 12.5 µL 2× master mix, 2.0 µL forward primer mix (0.5 µL per gene), 2.0 µL reverse primer mix (0.5 µL per gene), 2.0 µL cDNA template and 6.5 µL nuclease-free water. All samples were run in triplicate. Cycling was performed on a CFX96 system using: 95 °C for 2 min, followed by 40–45 cycles of 95 °C for 10 s and 58 °C for 30–60 s with data acquisition on FAM, HEX, Texas Red and CY5 channels. Each plate included non-template controls, water controls, and CY5-positive control wells to verify probe detection.

### 2.5 Data and Statistical Analysis

Relative gene expression analysis was performed using RT-qPCR targeting sdiA, csgD, flgG, fimA, rck, invA, bapA, and hilA, with 16S rRNA used as the internal housekeeping control. Raw fluorescence data were acquired over 36 amplification cycles and exported for downstream processing. Amplification curves were visually inspected to confirm sigmoidal behaviour and appropriate baseline characteristics. Ct values were determined using a fixed fluorescence threshold approach, whereby a global threshold of 500 RFU was applied uniformly across all reactions to ensure the threshold lay above baseline noise and within the exponential phase of amplification. Ct was defined as the first cycle at which RFU was ≥ 500. Target gene expression was normalised to the housekeeping gene to generate ΔCt values (Ct_target − Ct_16S), and treatment-induced changes were calculated relative to the LB control using the ΔΔCt method (ΔCt_treatment − ΔCt_LB). Relative expression was expressed as fold change using the equation 2−ΔΔCt and subsequently log₂-transformed for visualisation. Heat maps were generated to compare transcriptional responses across treatments and isolates. All data processing, statistical calculations, and graphical visualisations were performed using GraphPad Prism.

### 2.6 Molecular docking

Molecular docking analysis was performed to investigate potential interactions between quorum sensing molecules, quorum quenching compounds, and the *Salmonella Typhimurium* 14028 SdiA receptor. The study aimed to assess ligand-binding behaviour within the SdiA ligand-binding pocket and to identify compounds with potential quorum sensing inhibitory activity. The workflow incorporated protein sequence analysis, homology modelling, binding pocket prediction, ligand preparation, molecular docking, and structural visualisation.

The amino acid sequence of the *Salmonella Typhimurium* 14028 SdiA receptor was obtained from the publicly available whole genome sequence database (NCBI accession NZ_CP043907.1). Sequence comparison was performed against the experimentally characterised *Escherichia coli* SdiA receptor using UniProt and BLASTp analysis. Sequence alignment demonstrated that the *Salmonella* SdiA receptor shared approximately 71% sequence identity with the *E. coli* K-12 SdiA protein while maintaining identical protein length (240 amino acids), supporting the suitability of homology modelling using available *E. coli* SdiA crystal structures as templates. Conserved LuxR-family ligand-binding domains and DNA-binding domains were identified through InterProScan and Pfam analysis. Three-dimensional structural modelling of the *Salmonella* SdiA receptor was performed using SWISS-MODEL. The crystal structure of *E. coli* SdiA bound to 3-oxo-C6-homoserine lactone (PDB ID: 4Y15) was selected as the primary modelling template due to its ligand-bound conformation and high sequence similarity to the *Salmonella* receptor. Additional structural references included PDB structures 4LGW, 4Y13, and 2AVX. The generated homology model demonstrated acceptable structural quality metrics and preservation of the N-terminal ligand-binding domain associated with LuxR-family quorum sensing receptors. Binding pocket identification and validation were subsequently performed using ProteinsPlus/DoGSiteScorer and CASTp analysis. The primary predicted ligand-binding cavity exhibited a depth of approximately 10.06 Å and a volume of approximately 185.86 Å³, consistent with known LuxR-family receptor ligand pockets. Hydrophobicity analysis indicated a predominantly hydrophobic environment with hydrogen-bond donor and acceptor residues capable of stabilising acyl-homoserine lactone ligands. Ligand-binding coordinates were further validated by structural alignment of the generated *Salmonella* SdiA model against the experimentally resolved *E. coli* SdiA–3-oxo-C6-HSL complex (PDB: 4Y15) using PyMOL structural alignment tools. The coordinates of the bound ligand within the 4Y15 structure were extracted and used to define the docking search space for subsequent molecular docking simulations. Prior to docking, receptor structures were cleaned through removal of water molecules, solvent residues, inorganic ions, and non-essential chains using PyMOL. A focused ligand library was assembled to investigate native quorum sensing ligands alongside potential quorum quenching and anti-biofilm compounds. Native acyl-homoserine lactones included C4-HSL, C6-HSL, C8-HSL, and C12-HSL. Additional test compounds included farnesol, brominated furanone, cinnamaldehyde, thymoquinone, aloe-emodin, epigallocatechin gallate (EGCG), curcumin, allicin, and chlorhexidine. Ligand structures were obtained from the PubChem database in SDF format and prepared for docking using OpenBabel and SwissDock preparation tools. Molecular docking simulations were carried out using AutoDock Vina through the SwissDock platform. Docking calculations were centred around the predicted SdiA ligand-binding cavity using the previously validated coordinate set derived from the aligned 4Y15 structure. Each ligand was docked individually against the prepared *Salmonella* SdiA receptor model, and multiple docking poses were generated. Binding affinity values (kcal/mol) were recorded for each ligand, and the top-ranked docking conformations were selected for structural interpretation and comparative analysis. Docking visualisation and interaction analysis were performed using PyMOL. Ligand–receptor complexes were analysed to identify interacting residues located within 3.5–4 Å of each docked ligand. Hydrogen bonding interactions, hydrophobic contacts, and pocket occupancy were evaluated through residue selection, distance mapping, and surface rendering tools. The receptor was displayed using cartoon representations, while ligands and interacting residues were visualised using stick models and labelled according to residue identity and position. Structural surfaces surrounding the ligand-binding pocket were rendered to evaluate ligand accommodation within the cavity.

C8-HSL was used as the primary positive control ligand due to its structural compatibility with canonical LuxR-family quorum sensing receptors. Docking results demonstrated that C8-HSL occupied a deep hydrophobic cavity within the SdiA receptor and produced one of the strongest binding affinities observed among the tested ligands. Key interacting residues included Leu59, Val68, Val82, Leu83, Phe77, Phe100, Leu115, Tyr63, Tyr71, and Trp67, alongside polar residues such as Thr61, Gln72, Asp80, and Ser134 associated with stabilisation of the homoserine lactone headgroup. Comparative docking analysis demonstrated that shorter-chain ligands such as C4-HSL and C6-HSL retained similar binding orientations but exhibited reduced binding affinities, whereas the longer-chain C12-HSL demonstrated reduced accommodation within the pocket likely due to steric constraints.

Among the natural compounds tested, farnesol and thymoquinone demonstrated comparatively strong binding affinities and occupied the canonical ligand-binding pocket with interaction profiles overlapping those observed for the native AHL ligands. Aloe-emodin, cinnamaldehyde, and brominated furanone also docked within the same functional cavity, suggesting potential competitive binding behaviour. In contrast, EGCG, curcumin, and chlorhexidine produced weak or energetically unfavourable docking scores, indicating poor compatibility with the predominantly hydrophobic SdiA binding pocket. These findings were interpreted alongside phenotypic crystal violet assays and multiplex RT-qPCR gene expression analysis to assess potential quorum quenching activity and disruption of quorum sensing-mediated biofilm formation.

Homology modelling and molecular docking workflows were adapted from established SWISS-MODEL, SwissDock/AutoDock Vina, and PyMOL-based structural biology approaches. Waterhouse *et al*. (2018) for SWISS-MODEL; Grosdidier *et al*. (2011) for SwissDock; Trott and Olson (2010) for AutoDock Vina; DeLano (2002) for PyMOL

## 3. Results

### 3.1 CV biofilm Quantification (ST14028 and SE)

### 3.2 PCR and RT-PCR

**Figure 1.**
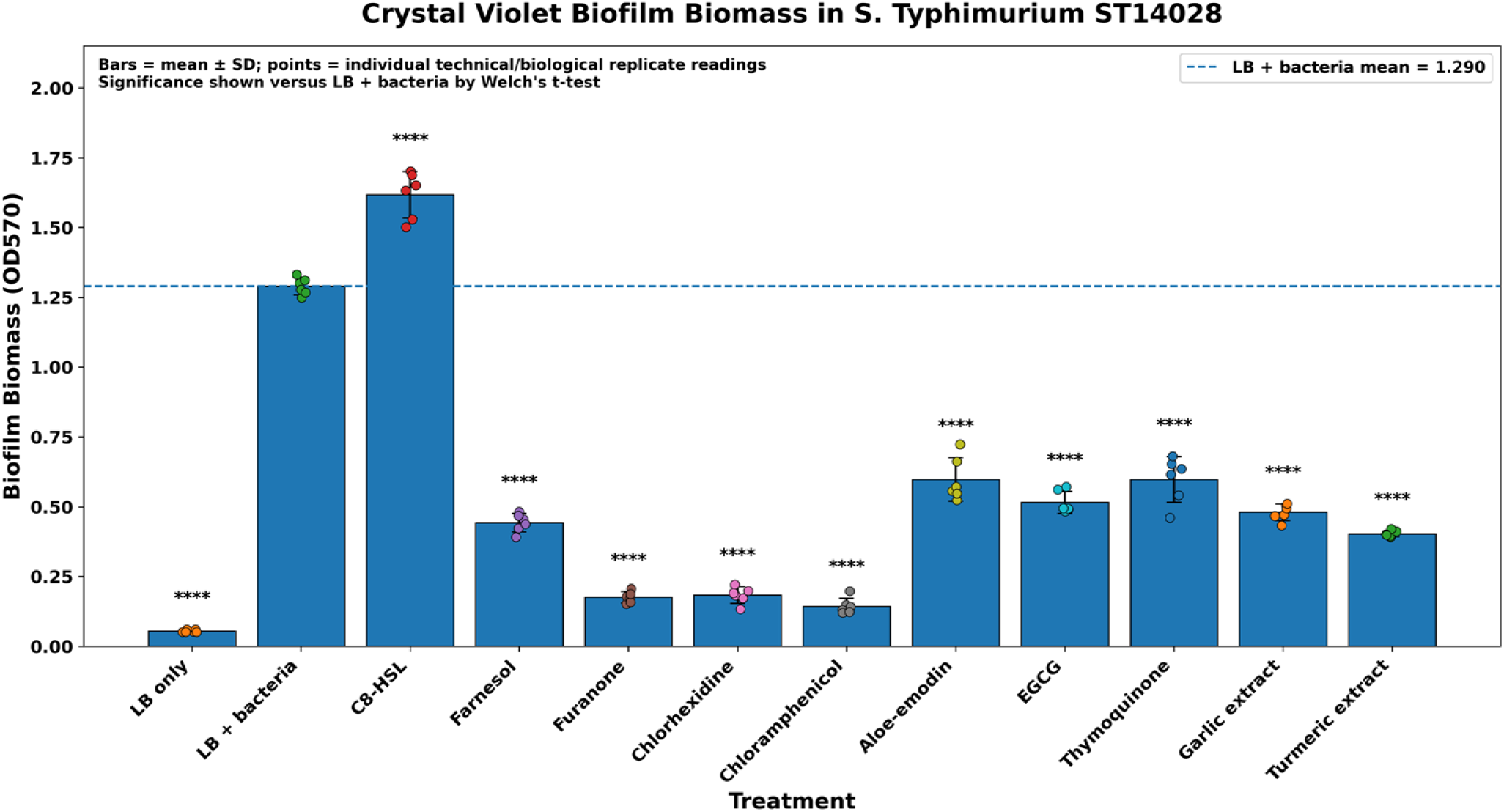

**Figure 2.**
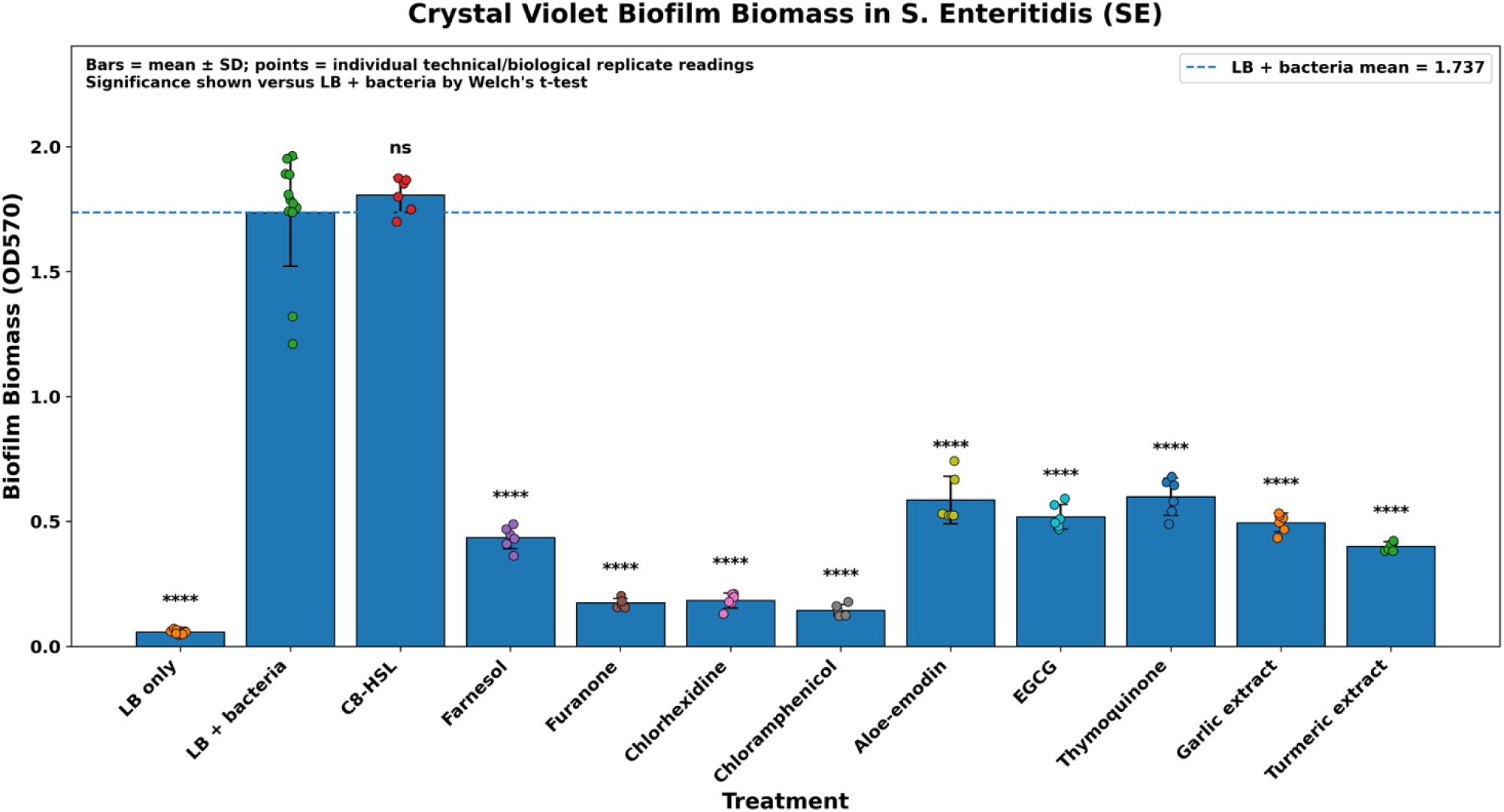

#### Figure 1 and 2 analysis

Crystal violet staining showed clear treatment-dependent changes in biofilm biomass in both *S. Typhimurium* ST14028 and *S. Enteritidis* SE. The LB-only control produced minimal OD570 signal, while the LB + bacteria control showed strong biofilm formation in both strains. In ST14028, C8-HSL significantly increased biofilm biomass compared with the LB + bacteria control, supporting a biofilm-enhancing effect of exogenous AHL stimulation. In contrast, farnesol, furanone, chlorhexidine and chloramphenicol significantly reduced biofilm biomass. The natural products, including EGCG, thymoquinone, aloe-emodin, garlic extract and turmeric extract, also reduced biofilm formation to varying degrees. A similar reduction in biofilm biomass was observed in SE following treatment with quorum-quenching agents, antimicrobial controls and natural products. However, unlike ST14028, C8-HSL did not significantly increase SE biofilm biomass, suggesting a strain-dependent response to exogenous quorum-sensing stimulation. Overall, these findings indicate that quorum-sensing and quorum-modulatory treatments differentially influence *Salmonella* biofilm formation in a strain-dependent manner.

**Figure 3.**
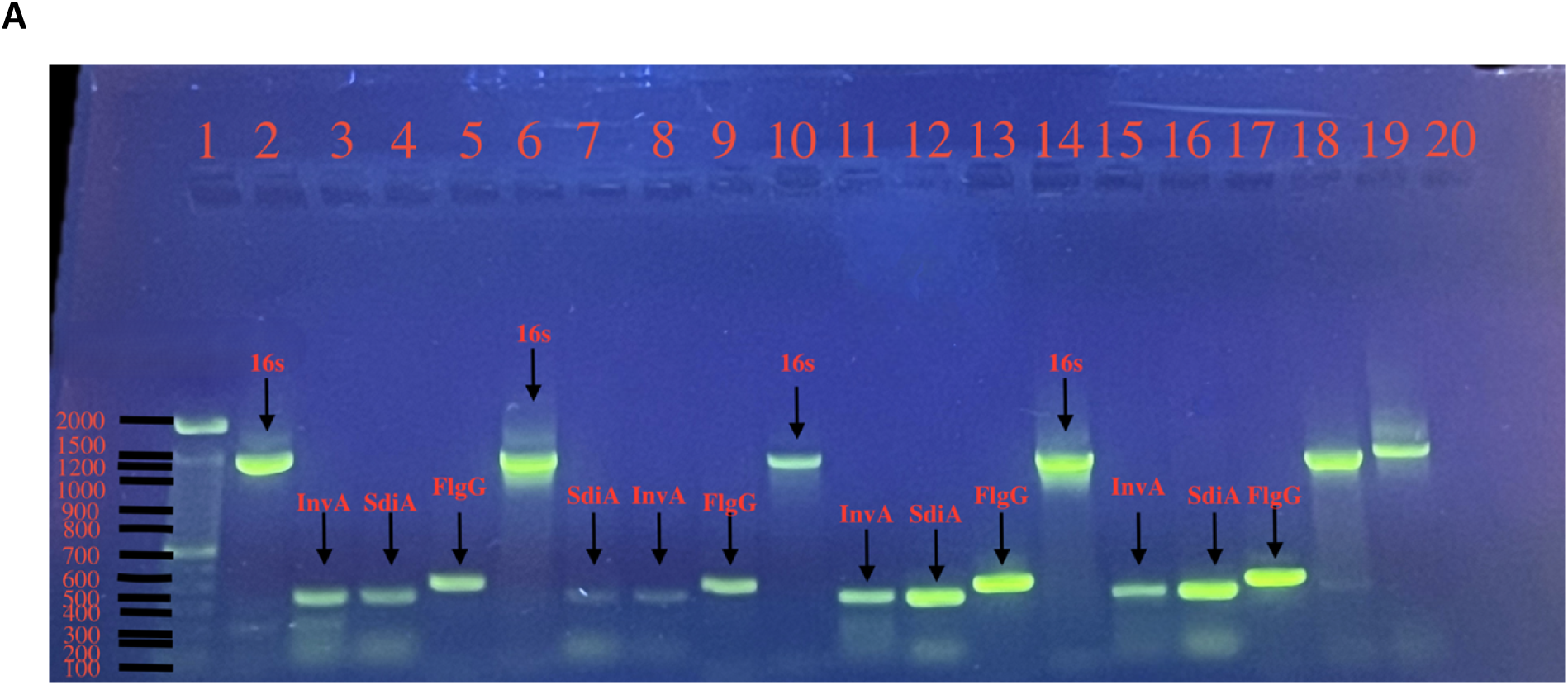
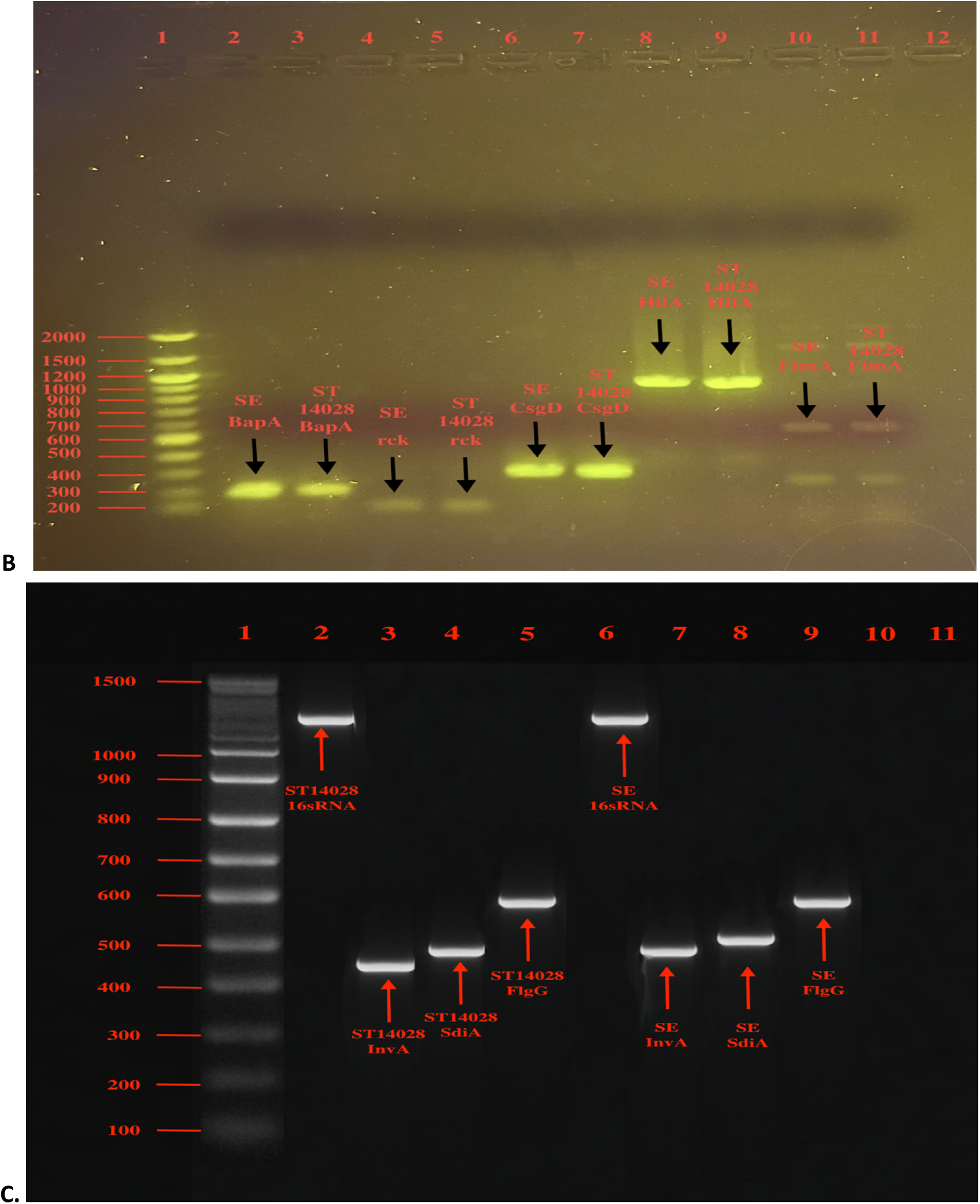
end point PCR gel images A, B and C.

**Figure 3**- Image A- Lanes 2 and 6 show PCR amplicons from S. Typhimurium 14028 DNA amplified with universal bacterial *16S rRNA* primers (∼1400 bp). Lanes 10 and 14 show similar PCR bands from *Salmonella* Enteritidis DNA templates, confirming the presence of the *16S rRNA* gene in all samples, which serves as a housekeeping gene. In Lanes 3 and 8 PCR amplicons from S. Typhimurium 14028 DNA amplified with the *Inv-a* gene (molecular weight 450 bp). lanes 11 and 15 show similar PCR bands from *Salmonella* Enteritidits DNA template. Lanes 4 and 7 contain S. Typhimurium 14028 DNA amplified with the *SdiA* gene primer, while lanes 12 and 16 contain S. Enteritidis DNA amplified with the same primer (500bp). Lanes 5 and 9 contain S. Typhimurium 14028 DNA amplified with the *FlgG* primer (∼600 bp), and lanes 13 and 17 contain S. Enteritidis DNA amplified with the same primer. Lanes 18 and 19 served as positive controls, using *E. coli* DNA amplified with the 16S rRNA primer. Lane 20 was a negative control containing no DNA template.

Image B - Lane 2 contains S. Enteritidis DNA amplified with the *BapA* primer (∼300 bp), and lane 3 contains S. Typhimurium 14028 DNA amplified with the same primer. Lane 4 contains S. Enteritidis DNA amplified with the *rck* primer, and lane 5 contains S. Typhimurium 14028 DNA amplified with the same primer. Lanes 6 and 7 contain S. Enteritidis and S. Typhimurium 14028 DNA, respectively, amplified with the *CsgD* primer (∼450 bp). Lanes 8 and 9 contain S. Enteritidis and S. Typhimurium 14028 DNA amplified with the *HilA* primer, while lanes 10 and 11 contain the same DNA order amplified with the *FimA* primer. Lane 12 served as a negative control containing no DNA template.

**Figure 4.**
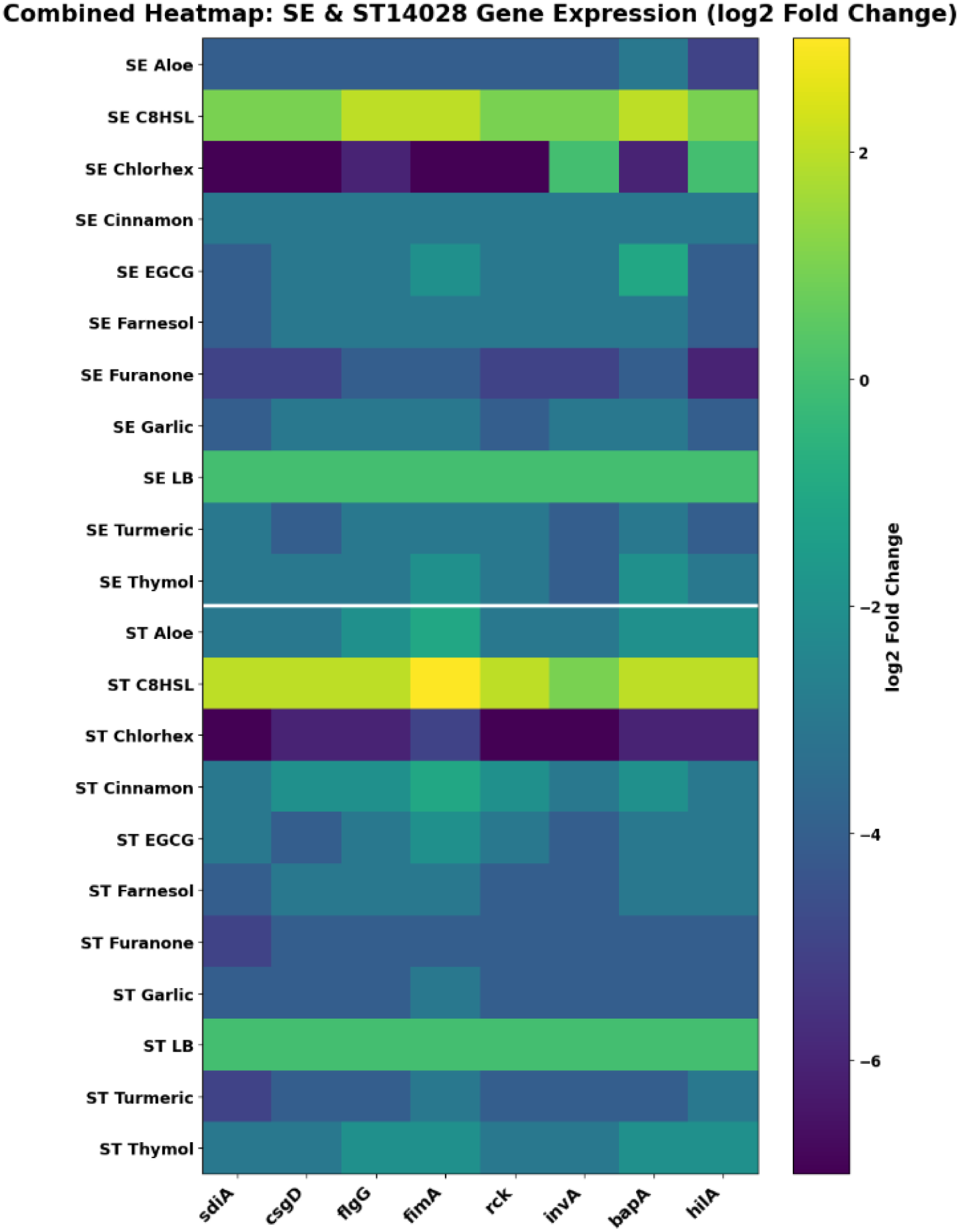
Relative Gene Expression (log₂ Fold Change) Determined by the ΔΔCt Method.

##### Figure 4 analysis

Combined heat map of relative gene expression in *Salmonella Enteritidis* (SE) and *S. Typhimurium* ST14028 following treatment with quorum-sensing molecules and phytochemical agents. Expression levels of *sdiA, csgD, flgG, fimA, rck, invA, bapA* and *hilA* were quantified by RT-qPCR and calculated using the ΔΔCt method, normalised to 16S rRNA. Values are presented as log₂ fold change relative to the LB control. Positive values indicate upregulation, whereas negative values indicate downregulation. In both strains, C8-HSL treatment produced broad upregulation of biofilm- and virulence-associated genes, while chlorhexidine caused marked transcriptional repression. Furanone and several phytochemical treatments also resulted in varying degrees of gene suppression, with clear strain-dependent differences observed between SE and ST14028.

### 3.3 3D Molecular docking binding affinity scores

**Figure 5.**
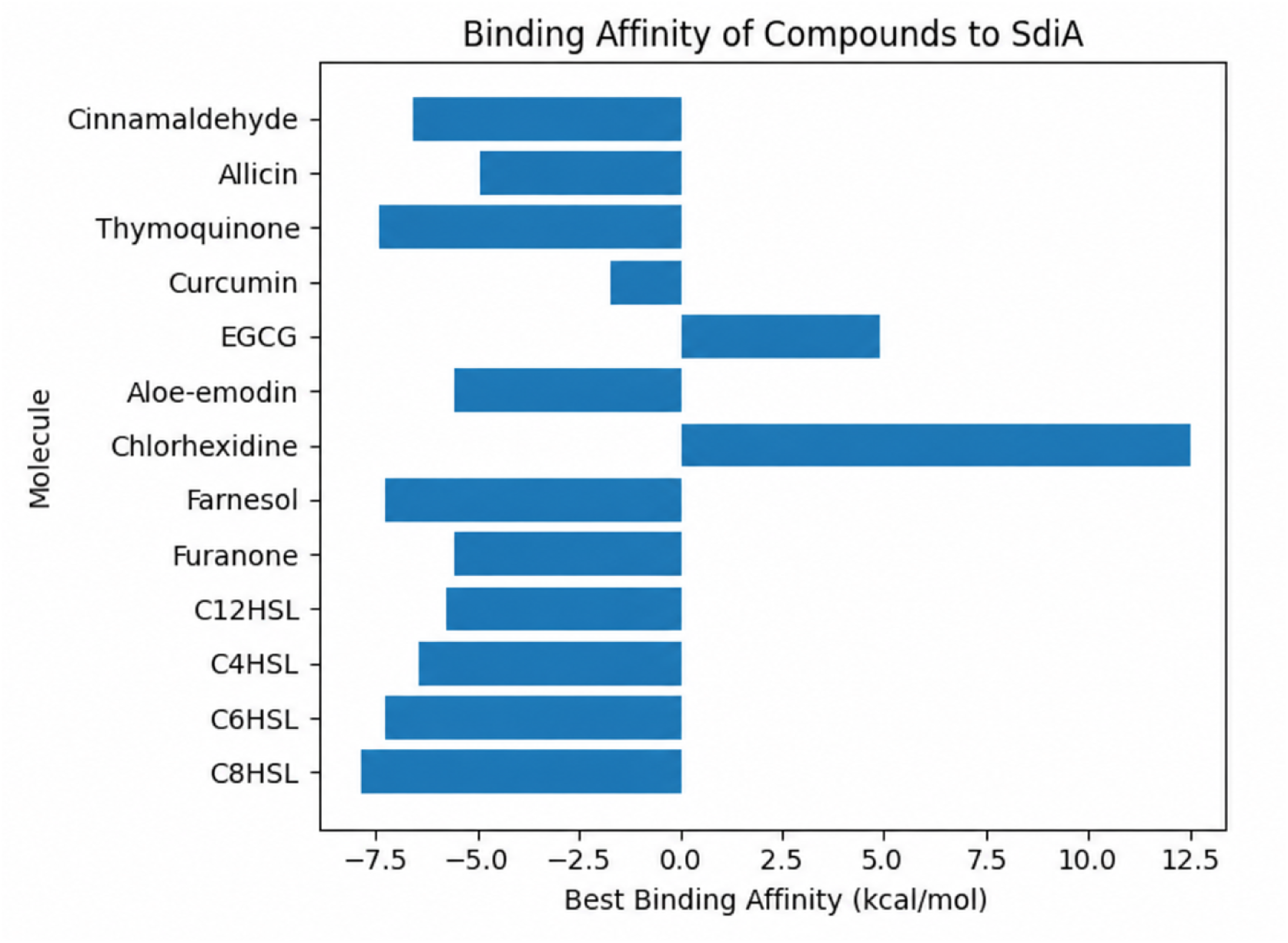

#### Figure 5 analysis

Molecular docking was used to assess the predicted interaction of quorum-sensing signals, quorum-quenching agents and selected phytochemical compounds with the *Salmonella* SdiA binding region. The AHL molecules showed favourable negative binding affinity values, with C8-HSL displaying strong predicted interaction with SdiA, supporting its role as an exogenous quorum-sensing signal. Farnesol and furanone also showed favourable predicted binding, consistent with their observed antibiofilm and transcriptional repression effects. Several phytochemical-associated compounds, including thymoquinone, aloe-emodin, allicin and cinnamaldehyde, also displayed negative binding affinities, suggesting possible interaction with SdiA. In contrast, EGCG and chlorhexidine showed weaker or unfavourable predicted binding, indicating that their antibiofilm effects may occur through mechanisms other than direct SdiA interaction.

## 4. Discussion

Biofilm formation by *Salmonella* enterica is a major determinant of its persistence on both abiotic and biotic surfaces, including food-contact materials and host gastrointestinal tissues (Lamichhane et al., 2024; Małaszczuk et al., 2025). Biofilm-associated cells exhibit increased tolerance to environmental stress, host immune responses, and antimicrobial treatments, complicating eradication efforts. The transition from planktonic growth to mature biofilm is tightly regulated at the genetic level by coordinating networks controlling adhesion, matrix production, motility, and virulence.The present study examined the transcriptional regulation of biofilm formation in S. Enteritidis (SE) and S. Typhimurium 14028 (ST14028) following exposure to quorum sensing (QS) molecules and the quorum quenching (QQ) agent farnesol. Expression profiling of eight target genes (sdiA, csgD, bapA, fimA, rck, hilA, invA, flgG) provided a multi-layered view of QS-mediated regulatory responses. As expected, reductions in cycle threshold (Ct) values corresponded to earlier or stronger transcriptional activation of these pathways.

### 4.1 Influence of AHLs on Gene Expression

Although *Salmonella enterica* does not synthesise acyl-homoserine lactones (AHLs) because it lacks a LuxI-type AHL synthase, it retains the LuxR homolog SdiA, which enables detection of exogenous AHLs produced by neighbouring microbial species. This ability allows *Salmonella* to respond to interspecies chemical cues within polymicrobial environments, including food-associated and host-associated biofilms. In the present study, C8-HSL was used as an exogenous quorum-sensing signal to determine whether external AHL exposure could reshape the expression of genes associated with biofilm formation, adhesion, motility and invasion in *S. Enteritidis* and *S. Typhimurium* ST14028.

Exposure to C8-HSL produced coordinated transcriptional activation across multiple QS- and biofilm-associated genes, including *sdiA, csgD, flgG, fimA, rck, invA, bapA* and *hilA*. This response was accompanied by increased biofilm biomass in the crystal violet assay, supporting the interpretation that external AHL sensing contributes to enhanced biofilm-associated phenotypes rather than reflecting a purely growth-related effect. The upregulation of *sdiA* suggests activation of the AHL-responsive regulatory pathway, while increased expression of *csgD* and *bapA* indicates stimulation of matrix-associated biofilm programmes. Similarly, changes in *fimA* and *flgG* expression suggest that adhesion and motility-related pathways are also influenced by exogenous signalling cues. The induction of *rck, invA* and *hilA* further indicates that AHL exposure may influence genes linked to host interaction and invasion-associated regulatory networks.

Importantly, the transcriptional response to C8-HSL differed between *S. Enteritidis* and *S. Typhimurium* ST14028, indicating strain-dependent sensitivity to quorum-sensing stimulation. This variation may reflect differences in baseline regulatory architecture, biofilm-forming capacity, or serovar-specific responses to environmental signalling molecules. Together, these findings support the model that *Salmonella* can exploit exogenous AHLs as interspecies cues to modulate biofilm-associated transcriptional programmes. Rather than acting as an autonomous AHL-producing organism, *Salmonella* appears to function as an AHL signal responder, using SdiA-mediated detection to adjust expression of genes involved in persistence, surface attachment, motility and virulence-associated behaviour. This reinforces the relevance of SdiA-mediated interspecies signalling as a regulatory layer in *Salmonella* biofilm biology and provides a mechanistic basis for evaluating quorum-quenching and phytochemical compounds as potential modulators of these responses.

### 4.2 Quorum Quenching Effects of Farnesol

Farnesol has been widely reported to disrupt biofilm development across multiple microbial species. In the present study, farnesol exposure produced consistently higher Ct values for several biofilm-associated genes, indicating delayed or reduced transcriptional activation. The most pronounced suppression was observed for csgD and bapA, suggesting that farnesol primarily interferes with matrix regulatory pathways rather than exerting a purely bactericidal effect.

These transcriptional patterns support a quorum quenching mechanism in which farnesol disrupts QS-responsive signalling upstream of biofilm structural gene expression. Importantly, these genotypic changes corresponded with reduced biofilm biomass observed phenotypically, strengthening the functional relevance of the transcriptional data.

While prior studies have demonstrated the antibiofilm activity of farnesol in mixed microbial systems, the present findings extend this evidence by demonstrating direct modulation of QS-linked gene expression in Salmonella. This supports the concept that quorum quenching agents may attenuate pathogenic behaviours without necessarily inhibiting planktonic growth.

### 4.3 Integration of Phenotypic and Genotypic Outcomes

A key strength of this study is the integration of biofilm quantification with targeted transcriptional analysis. Previous crystal violet assays demonstrated that AHL supplementation enhanced biofilm biomass without significantly altering planktonic growth kinetics. The RT-PCR data presented here provide mechanistic support for this observation, showing that QS stimulation accelerates activation of adhesion, matrix, and virulence-associated genes.

Conversely, farnesol treatment delayed transcriptional activation across multiple nodes of the QS–biofilm network. The coordinated response observed across rck, fimbrial genes, and SPI-1–associated regulators further supports the existence of a multi-gene regulatory axis linking quorum sensing to Salmonella pathogenic behaviour.

Collectively, the combined phenotypic and genotypic data strongly support the conclusion that QS-mediated transcriptional reprogramming contributes directly to enhanced biofilm formation in Salmonella enterica.

## 5. Conclusion

This study demonstrates that quorum sensing (QS) signalling, particularly via short- to medium-chain AHLs, accelerates the transcriptional activation of key biofilm- and virulence-associated genes in *Salmonella enterica*. RT-PCR analysis revealed consistently lower Ct values for **sdiA,** csgD, bapA, fimA, rck, hilA, invA, and flgG in AHL-treated samples, indicating earlier onset of gene expression compared with untreated controls. These transcriptional changes align with increased biofilm biomass observed phenotypically, supporting the role of QS as a regulatory driver of *Salmonella* biofilm development rather than a simple growth effect.

In contrast, the quorum quenching agent farnesol produced delayed transcription across multiple biofilm-associated genes, particularly csgD and bapA, consistent with suppression of matrix regulatory pathways. These findings suggest that quorum interference strategies may attenuate biofilm formation by disrupting coordinated gene activation.

To our knowledge, this study provides one of the more integrated transcriptional assessments of QS-mediated biofilm regulation in *S. Enteritidis* and *S. Typhimurium* using a multi-gene panel. The results highlight the potential of targeting QS-regulated networks as an anti-virulence approach for controlling *Salmonella* persistence in clinical and food-associated settings.

Future work should incorporate broader transcriptomic approaches (e.g., RNA-seq), functional mutagenesis, and in situ biofilm models to further delineate the QS regulatory landscape and to validate candidate quorum-modulatory compounds.

## Data Availability Statement

All relevant data generated or analysed during this study are included in the manuscript. Raw data files, including OD readings for CV assay and multiplex raw data are available here https://doi.org/10.5281/zenodo.20382743

## Ethics Statement

The study did not involve human participants, animal models, or patient data.

## Acknowledgement

This manuscript used a Large language model (Chatgpt version 4.0) for language correction.

## Disclosure statement

The authors report there are no competing interests to declare.

Biographical note for Schnieder Fernandes: I am a second-year PhD student conducting research on bacterial genomes with a focus on Salmonella serovars. My project involves using CRISPR-Cas9 technology to generate mutant libraries, allowing me to investigate the functions of unknown genes. With a background in cell and molecular biology as well as immunology, I am currently developing a plasmid-free gene editing strategy for this research.

Biographical note for Anjali Ghosh: My primary research focus for the past twenty-five years has been on bacterial molecular biology, particularly the pathogenesis of *Salmonella enterica* serovars. During this time, I have investigated key aspects of bacterial behaviour, including multidrug resistance (MDR), virulence, biofilm formation, and overall pathogenesis, contributing to a deeper understanding of bacterial infections and their implications for therapeutic interventions. These works have been published in peer-reviewed journals. In addition to my work in bacterial molecular biology, I have also researched in immunology, focusing on ACE gene polymorphism, specifically exploring their roles in conditions such as angioedema and food and venom allergies. My earlier investigations into the effects of plant oestrogens on breast cancer cell lines also yielded significant findings, which were published in respected scientific journals.

